# Predictors of genomic differentiation within a hybrid taxon

**DOI:** 10.1101/2021.04.11.439335

**Authors:** Angelica Cuevas, Fabrice Eroukhmanoff, Mark Ravinet, Glenn-Peter Særtre, Anna Runemark

**Affiliations:** Department of Biosciences, Centre for Ecological and Evolutionary Synthesis, University of Oslo, PO Box 1066, N-0316, Oslo, Norway; School of Life Sciences, University of Nottingham, NG7 2RD, Nottingham, United Kingdom; Department of Biology, Lund University, Sölvegatan 37, SE-22362, Lund, Sweden

## Abstract

Hybridization is increasingly recognized as an important evolutionary force. Novel genetic methods now enable us to address how the genomes of parental species are combined to build hybrid genomes. However, we still do not know the relative importance of contingencies, genome architecture and local selection in shaping hybrid genomes. Here, we take advantage of the genetically divergent island populations of Italian sparrow on Crete, Corsica and Sicily to investigate the predictors of genomic variation within a hybrid taxon. We test if differentiation is affected by recombination rate, selection, or variation in ancestry proportion from each parent species. We find that the relationship between recombination rate and differentiation is less pronounced within hybrid lineages than between the parent species, as expected if purging of minor parent ancestry in low recombination regions reduces the variation available for differentiation. In addition, we find that differentiation between islands is correlated with differences in selection in two out of three comparisons. Patterns of within-island selection are correlated across all islands, suggesting that shared selection may mould genomic differentiation. The best predictor of strong differentiation within islands is the degree of differentiation from house sparrow, and hence loci with Spanish sparrow ancestry may vary more freely. Jointly, this suggests that constraints and selection interact in shaping the genomic landscape of differentiation in this hybrid species.

## INTRODUCTION

Heritable variation is the substrate on which natural selection acts, and hybridization is increasingly recognized as an important process providing such variation in fish [1, 2], insects [3], birds [4, 5] and even humans [6]. Hybridization can enable species to combine parental genomes in adaptive ways, for instance contributing alleles linked to insecticide resistance in mosquitoes malaria vectors [7], adaptive fur colour in hares [8] and MHC immune defence diversity in modern humans [9]. Similarly, the variation created by hybridization has provided the raw materials for the extensive adaptive radiations of African lake cichlids [2,10,11,12,13]. Lineages resulting from hybridization may even outcompete the parental species in certain environments and colonize new niches as documented in *Helianthus* sun flowers where hybrid taxa colonize extreme environments [14, 15]. One outcome of hybridization is hybrid speciation, resulting in the formation of a taxon that is reproductively isolated from its parent species [16]. Hybrid speciation can arise both through allopolyploidization and homoploid hybrid speciation, without increase in chromosome number [16,17,18,19]. Interestingly, the relative contributions of the parental species may vary within a hybrid taxon, as illustrated by the variable genome composition in sword-tail guppies [20], in Lycaides butterflies [21], and in isolated island populations of Italian sparrows [22].

Hybridization, at least in animals, was historically viewed as an evolutionary mistake [23], partly because hybrids are likely to suffer from incompatible combinations of derived alleles. While this view has changed over the last decades [17, 24], hybrid lineages likely need to overcome a number of challenges to establish. Incompatibilities might mean low fertility, sterility or even inviability in some crosses [25]. This is shown by Haldane’s rule [26], when species have heterogametic sex chromosomes, the heterogametic sex is more likely to be sterile or inviable. In addition, evidence for a role of mito-nuclear interactions causing fitness reduction in hybrids is mounting [27,28,29]. For example, maladaptive metabolisms in hybrid species [30] suggest that mito-nuclear interactions could pose strong selection pressures on the genomic composition in hybrid taxa. Mito-nuclear interactions may also play a role in determining Italian sparrow genome composition [27]; hybrid Italian populations are largely fixed for house sparrow mitochondrial genomes, and there is evidence of an excess of house sparrow ancestry conserved in nuclear genome regions contributing to mitochondrion function [22]. Even in species that have successfully formed hybrid daughter lineages, early generation hybrids may still be inviable or infertile [31]. These findings suggest that only specific combinations may be viable in a hybrid lineage or only specific portions of the genome are free to vary, potentially resulting in convergent genomic compositions of hybrid lineages.

Although hybrid lineages in principle have a vast number of potential combinations of parental alleles and increased nucleotide diversity available as a source for adaptation, little is known about genome stabilization in hybrid taxa [19, 32]. There could potentially be constraints on genomic mosaicism and bias on the overall composition of hybrid genomes. For instance, in nightingales [33], flycatchers [34] and field crickets [35] introgression on the sex chromosomes was reduced compared to genome-wide levels, consistent with selection against infertility. Experimental assays in sunflowers have shown that the same genetic combinations found in natural hybrid lineages re-emerge in experimental hybrid populations [36], possibly due to selection against alternative combinations. This raises the question of how easily hybrid lineages can achieve divergent genome compositions and phenotypes. Can different combinations of parental alleles easily be achieved due to selection for divergent local adaptation in homoploid hybrids? Or do patterns of differentiation at a local scale mirror those between strongly divergent populations, suggesting a role for constraints on which genome regions may differentiate? Exploiting the patterns of population differentiation within hybrid species may reveal novel insights into the selective forces shaping hybrid genomes.

Interestingly, patterns of species differentiation are affected by the recombination rate landscape [37,38,39]. This can result in highly correlated patterns of divergence between closely related species pairs, such as that found in flycatchers [37]. Moreover, linked selection is more efficient in low recombination regions [37]. Selective sweeps in genomic regions of low recombination can give rise to a negative correlation between recombination rate and genomic differentiation [37, 40]. Specific to hybrid taxa, purging of minor parent ancestry in low recombination regions to reduce genomic incompatibilities could reduce the variation available for differentiation. Recent studies have indeed found reduced introgression in low recombination regions in hybrid swordtail fish, sticklebacks, *Heliconius* butterflies and humans, suggesting that the recombination landscape may indeed affect which regions are permeable to introgression [20,38,41]. Genomic blocks from the minor-parent may be retained in regions of high recombination rate due to their increased likelihood of breaking away from potential incompatibilities [20]. If purging of incompatibilities in low recombination regions is pervasive, the relationship between recombination rate and differentiation is expected to be less negative in hybrid lineages compared to the differentiation between parental taxa (Fig. 1A), but the relative importance of these two processes in shaping hybrid genomes remains unclear.

**Figure 1.**
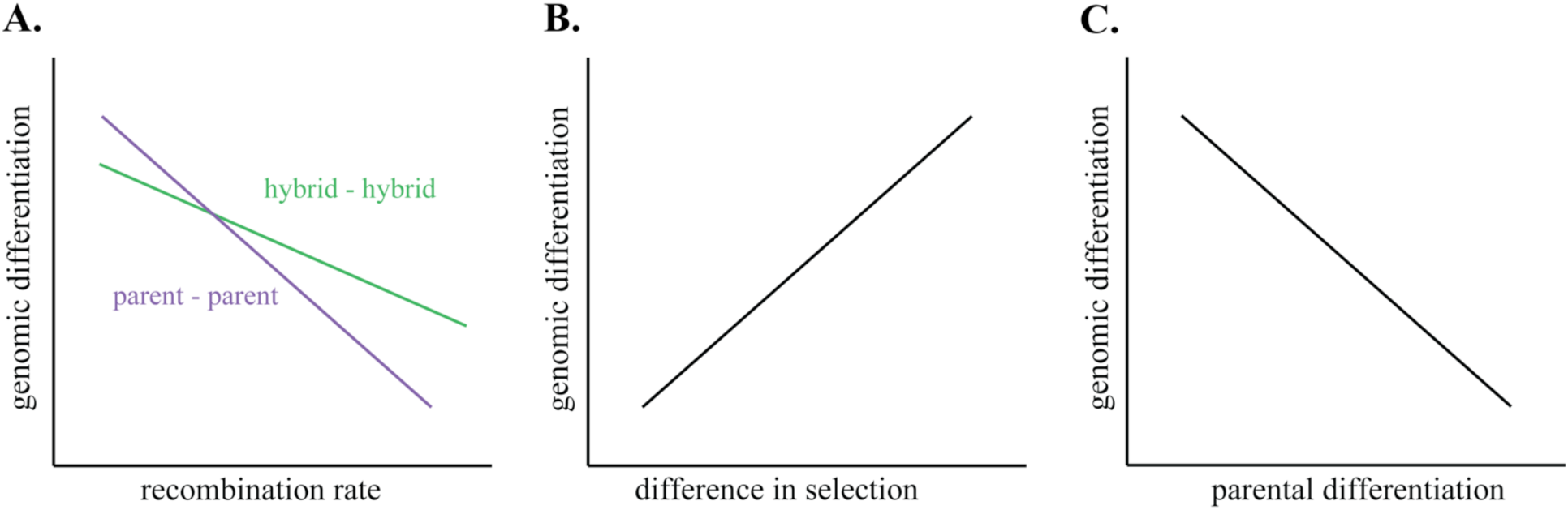
Expected predictors of within-island genomic differentiation. **A. Recombination rate:** *Parent-parent:* Since selective sweeps are more efficient in low recombination regions we expect a negative relationship between recombination rate and differentiation between the non-hybrid parental lineages. ***Hybrid - hybrid:*** Specific to hybrid taxa, additional selection against minor ancestry in low recombination regions could reduce variation and the potential for differentiation. As this process does not affect parent taxa, we would expect a flatter relationship between recombination rate and differentiation in hybrids, compared to their parent taxa, if purging of minor parent ancestry is pervasive. Also, in higher recombination regions there is a potential for alternative blocks of ancestry to be fixed, which will inflate the differentiation among hybrids as recombination rate increases, relative to the parent-parent comparison, contributing to the flatter relationship in hybrids **B. Selection:** If selection is an important predictor of differentiation, we expect that differentiation among populations is positively correlated to differences in selection among populations. **C. Parental differentiation:** If differentiation is limited by potential incompatibilities between divergent parental loci we would expect that the potential for adaptive differentiation in the hybrid would be highest in genomic regions of low parental differentiation. In contrast, highly differentiated regions, more likely to be under constraint due to incompatibilities between parents, would have a greater likelihood of being fixed for the ancestry of one parent only.

There is evidence that hybrid taxa can use the variation that originated through hybridization for local adaptation in the hybrid taxon. For instance, beak shape in Italian sparrows is explained by local precipitation regime [42]. Beak size differences between island populations are best explained by temperature seasonality [43] and the island populations are strongly differentiated for a gene known to affect beak morphology in Darwin’s Finches, FGF10 [22, 44]. In addition, the gene GDF5, part of the BMP gene family that has a fundamental role in beak shape and size variation in Darwin’s finches [45, 46], is a candidate gene putatively under selection in the Italian sparrow populations from mainland Italy [32]. However, beak shape is also affected by the proportion of the genome inherited from each parent species suggesting a small albeit significant role for contingencies in morphology [42, 43]. Moreover, in a recent study investigating genetic differentiation among Italian sparrow populations on mainland Italy, population differentiation was associated with temperature and precipitation [32]. Interestingly, alleles segregating in both parental species showed strong allele frequency differences within the Italian sparrow, suggesting that adaptation is not completely contingent on the combination of alleles from different parent species [32]. Selection, fixing alleles from either one or the other parent depending on which variant is favoured locally, could contribute to the stabilization of hybrid genomes. If divergent selection pressures between islands, favouring the fixation of alternate parental alleles, are important for differentiation, we would expect stronger differentiation among islands in regions under divergent selection (Fig 1B). However, genomic regions of parental divergence are more likely to harbour incompatibilities, thus fixation for ancestry of one parental allele only is likely to occur in the hybrid due to constraints. Consequently, local differentiation in the hybrid could be expected to be highest in genomic regions of low parental differentiation (Fig 1C).

One way to further our understanding of the selective forces acting on hybrid genomes is to investigate patterns of differentiation within hybrid lineages and the factors that best explain them. Here, we focus on the Italian sparrow, a clear example of a homoploid hybrid species, with reproductive barriers to the parent species consisting of a subset of those isolating the parent species [27, 47]. This is a uniquely suited study system as there are genetically divergent island populations, potentially originating from independent hybridization events, with different proportions of their genomes inherited from their parental species, house and Spanish sparrows (*P. domesticus* and *P. hispaniolensis*) [22] (Fig. 2). We use the island Italian sparrows to investigate how differentiation *among populations within each island* compares to *among island* differentiation. Our aim is to provide insight into how readily within-island local populations of hybrid species differentiate in relation to differentiation between islands. We also address which factors best predict differentiation, specifically addressing whether recombination rate, differences in selection or ancestry contribution best predict differentiation within- and among islands.

**Figure 2.**
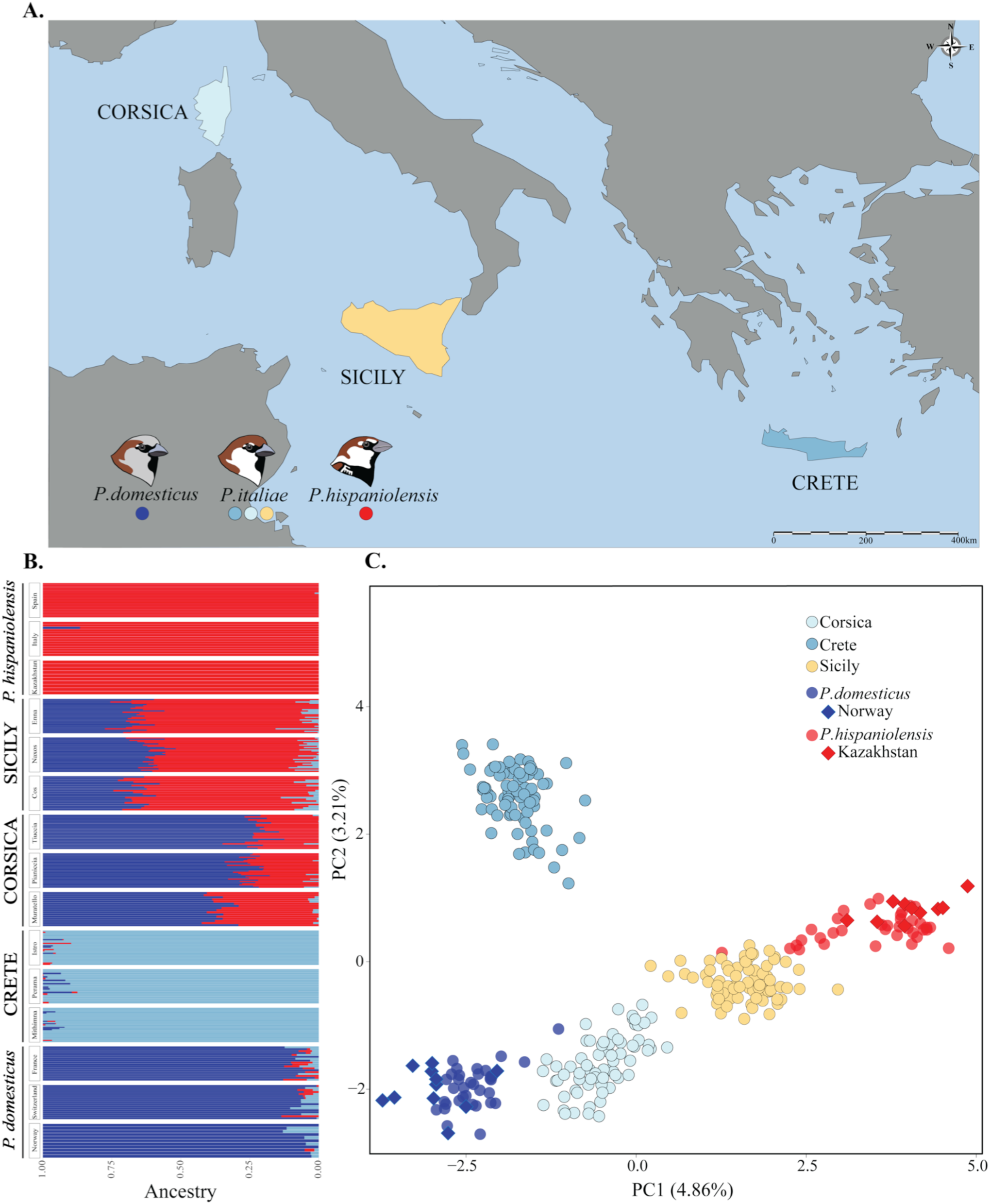
Sampling design and population structure. **A.** Map of sampling locations (map redrawn from Runemark et al., 2018a). Sparrows were sampled in three populations in each island, Corsica (N=70), Crete (N=77) and Sicily (N=76) **B.** Admixture analysis illustrating the clustering of the island populations of the Italian sparrow and their parental species for three clusters (k=3), the value with highest support. Three geographically separated populations of the parental species, the house sparrow and Spanish sparrow, were included. **C.** Principal component analysis illustrating the relationship between the Italian sparrow populations and the parental species. ♦ indicate the reference parental populations with lower levels of introgression.

We test the hypotheses that I) long periods of independent evolution have resulted in significantly higher divergence *among*-islands than *within*-islands; II) selective sweeps and purging of minor parent ancestry in low recombination regions has led to a correlation between recombination rate and differentiation that is lower than that between the parent species (Fig. 1A) III) that regions under stronger differential selection between islands also are more differentiated *among* islands (Fig. 1B) IV) that constraints on how freely genomic regions are able to diverge have led to a correlated landscape of *within*-island and *among*-island differentiation, and V) that differences in minor – major parental ancestry in the hybrid have a direct effect on the genomic differentiation between populations.

## RESULTS

### I) Genomic differentiation within- and between islands

Consistent with [22] we find strong differentiation between the studied islands. From our principal component analysis, the first main axis of differentiation largely reflects the proportion of the genome inherited from each parent species, and Crete diverges along the second axis (Fig. 2C). Interestingly, the ADMIXTURE analysis supports the presence of three clusters rather than two, with Crete forming a separate cluster (Fig. 2B). *F*_ST_ estimates are consistent with this, with Crete being overall more differentiated from Sicily (mean *F*_ST_ = 0.043) and Corsica (0.042), while mean *F*_ST_ between Corsica and Sicily is lower (0.025; Fig. 3A). *F*_ST_ estimates between islands are higher than those within islands (Monte-Carlo permutation paired t-test (t=33.21, df=7927, *P*=1.98e-15; Table S3; Fig. 3A). A discriminant function analysis recovers some differentiation among local populations in each island (Fig. 3B, Fig. 3C), and correctly assigns 95.3% of Corsican individuals, 78.4% of the individuals from Crete and 75.5% of the Sicilian individuals to their *within*-island populations of origin. Within-island *F*_ST_ differs significantly among islands (All Ps < 0.5e-3; Table S3), with Corsican populations exhibiting the highest mean *F*_ST_ of 0.018, as well as the highest nucleotide diversity (π: 3.021e-06; Table S3). Differentiation within Sicily is intermediate at 0.013, while Crete has the lowest within-island *F*_ST_ of 0.011 (Fig. 3A, Table S3). While most variation segregates within individuals and populations, an AMOVA reveals that 4.84% of the variation is found among islands whereas 0.91% of the variation is found among populations within islands (both fractions are statistically significant *P:* 0.001, as estimated from a randomization Monte Carlo test with 1000 permutations; Table 1, Fig. S1).

**Figure 3.**
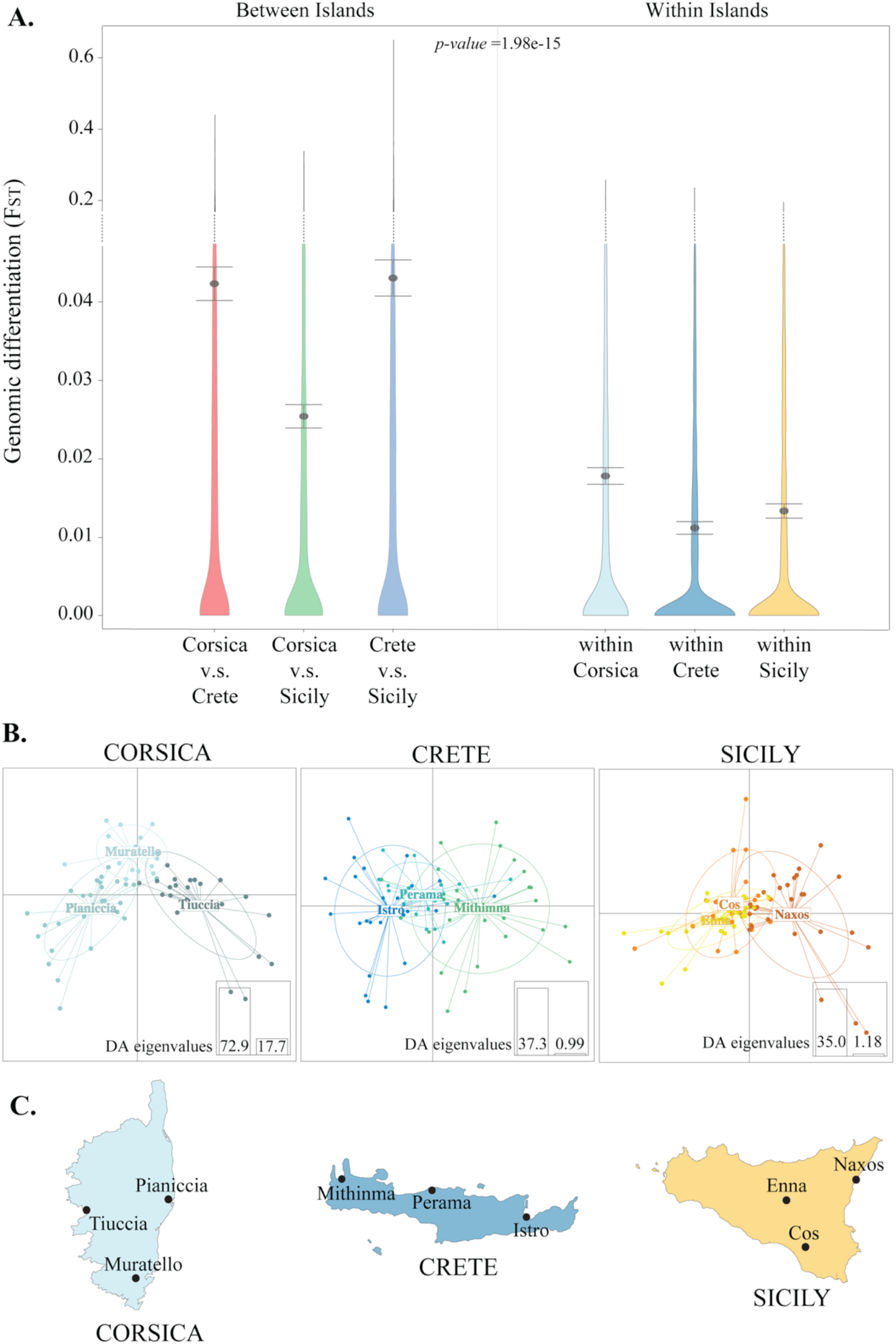
Genomic differentiation in islands. **A.** Levels of genomic differentiation *among-* islands compared to levels of differentiation between populations *within-*islands. Means and 95% CI depicted. **B.** Discriminant analysis of principal components (DAPC) for populations in Corsica, Crete and Sicily. **C**. Map of studied islands with the three populations sampled within each island.

**Table 1.** Analysis of Molecular Variance (AMOVA) across islands and populations within islands. Several cut-offs for missing-ness per loci were use (5%, 10%, 20%), but the results from the AMOVA did not chance substantially. Significance of variation partitioning elements was maintained.

| Analysis of Molecular Variance |  |  |  |  | Randomization by |  |
| --- | --- | --- | --- | --- | --- | --- |
| AMOVA |  |  |  |  | Permutation |  |
|  |  |  |  |  | Monte Carlo test |  |
| Variance partitioning | Df | Sum Sq | Sigma | % of covariance | Std. Observed | *P-value* |
| Among islands | 2 | 778.79 | 2.22 | 4.84 | 4.33 | 0.001** |
| Between populations within island | 6 | 365.37 | 0.42 | 0.91 | 9.42 | 0.001** |
| Between individuals within populations | 213 | 8654.88 | -2.54 | -5.54 | -4.17 | 1.000 |
| Within individuals | 222 | 10148.04 | 45.71 | 99.80 | 1.26 | 0.897 |

### II) The relationship between genomic differentiation and recombination rate

We tested the difference in the slopes of the relation between recombination rate and genomic differentiation for each island and the parental divergence. We also evaluated whether there was a significant interaction between comparison (e.g. parent-parent vs. within island) and recombination rate on differentiation for each island. This allows us to test the hypothesis from Fig 1A; i.e. we predicted that the slopes would be less steep in the hybrids if variation and hence differentiation is decreased in low recombination regions in hybrids due to selection against minor parent ancestry, and find that this is the case (Table S4; Table S5; Fig. 4A). The slope generated by the relation between the parental differentiation and recombination rate differs from those found in each within-island comparison (Table S4; Fig. 4A). We find a significant interaction of recombination rate and comparison (e.g. parent-parent vs. within island) in all independent linear models per island (Table S5). We find no significant correlation between recombination rate and within-island genomic differentiation for Corsica (correlation=-0.012, R^2^=1.4e-4, *P*=1; Fig. 4B) or Crete (correlation= - 0.003, R^2^=0.9e-5, *P*=1; Fig. 4C). However, differentiation within Sicily is weakly but significantly negatively correlated with recombination rate, (correlation=-0.048, R^2^=0.002, *P*=0.037; Fig. 4D).

**Figure 4.**
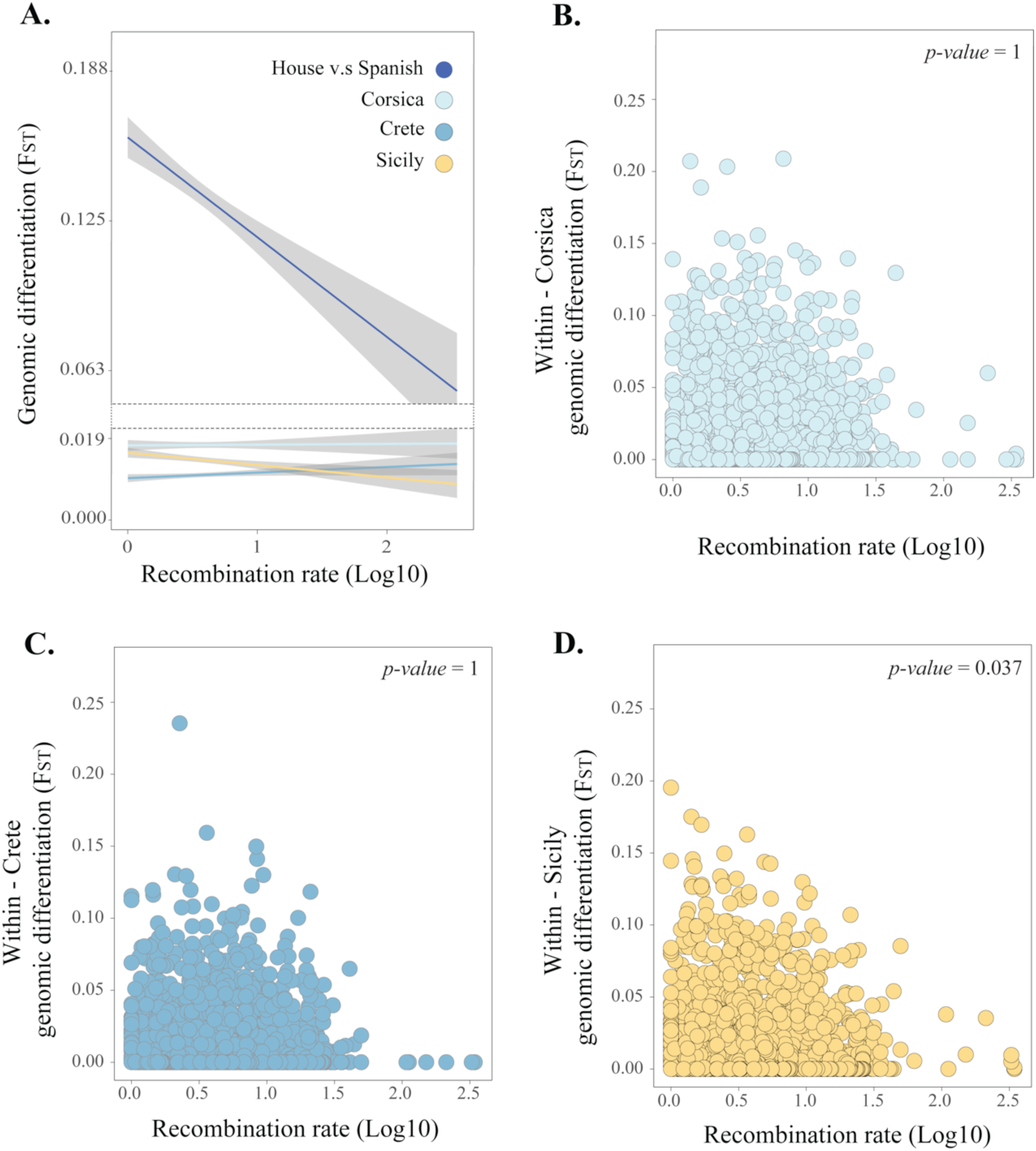
The influence of recombination rate on genomic differentiation. **A.** Comparison of the relationship between recombination rate and genomic differentiation between the parental species (dark blue) and between Italian sparrow populations within each island. Correlation between recombination rate and *within*-island genomic differentiation for **B**. Corsica, **C.** Crete and **D.** Sicily.

A GLM examining the influence of recombination rate, parental differentiation and differentiation to the two parent species, respectively, did not reveal any significant relationship between recombination rate and differentiation within islands (Estimate=-9.66e-04, Std.Error=7.84e-04, *P*=0.22; Table S7). A corresponding binomial model revealed that recombination rate did not affect the probability of loci being *F*_ST_ outliers within islands either (Table 2, Table S6). However, a corresponding GLM evaluating individual islands separately shows that recombination rate significantly explains differentiation between Sicilian populations (Estimate=-3.48e- 03,Std.Error=1.34e-03, *P*=9.3e-3; Table S8; Fig. 4D).

**Table 2.** Logistic regressions on the probability of being a local F_ST_ outlier within island.

| **Model:** |  |  |  |  |
| --- | --- | --- | --- | --- |
| Pr(outlier)= Hybrid Index + Recombination Rate + island + island.house $F_{ST}$ + island.Spanish $F_{ST}$ + house.Spanish $F_{ST}$ | | | | |
| **Response variable** | **Predictor** | **Estimate** | **Std. Error** | **P-value** |
| Pr (within-island $F_{ST}$ outlier) | Recombination Rate | -1.402e-01 | 1.431e-01 | 0.327 |
|  | Hybrid Index | 1.411e-02 | 6.336e-02 | 0.824 |
| | Island v.s House $F_{ST}$ | -1.401e+00 | 6.674e-01 | 0.036* |
| | Island v.s Spanish $F_{ST}$ | 5.728e-01 | 4.698e-01 | 0.223 |
| | House v.s Spanish $F_{ST}$ | -6.752e-06 | 8.770e-06 | 0.441 |

**Post-hoc Estimated marginal (Least-squares) means for the predictor variable “island”**
| **Comparison** | **Estimate** | **Std. Error** | **Z ratio** | **P-value** |
| --- | --- | --- | --- | --- |
| Corsica - Crete | 0.0207 | 0.133 | 0.159 | 0.9866 |
| Corsica - Sicily | -0.1135 | 0.133 | -0.877 | 0.6684 |
| Crete - Sicily | -0.1343 | 0.134 | -1.025 | 0.5756 |

*p-value* adjustment: Tukey

### III) The concordance of patterns of selection and genomic differentiation

To assess the role of selection in shaping genomic differentiation in the Italian sparrow, we test if differentiation is significantly correlated to differences in selection. We find that genomic differentiation between island pairs was significantly correlated to divergent selection. *F*_ST_ and xp- EHH were negatively correlated for the Corsica – Sicily (correlation = -0.061, R^2^=0.0037, *P* = 0.014), and Crete and Sicily (correlation = -0.059, R^2^=0.004, *P* = 0.019; Fig. 5A; Table S9A) comparisons. However, we do not find any significant correlation between differentiation and selection differences for the Corsica – Crete combination (correlation = -0.042, R^2^=0.002, *P* = 0.148).

**Figure 5.**
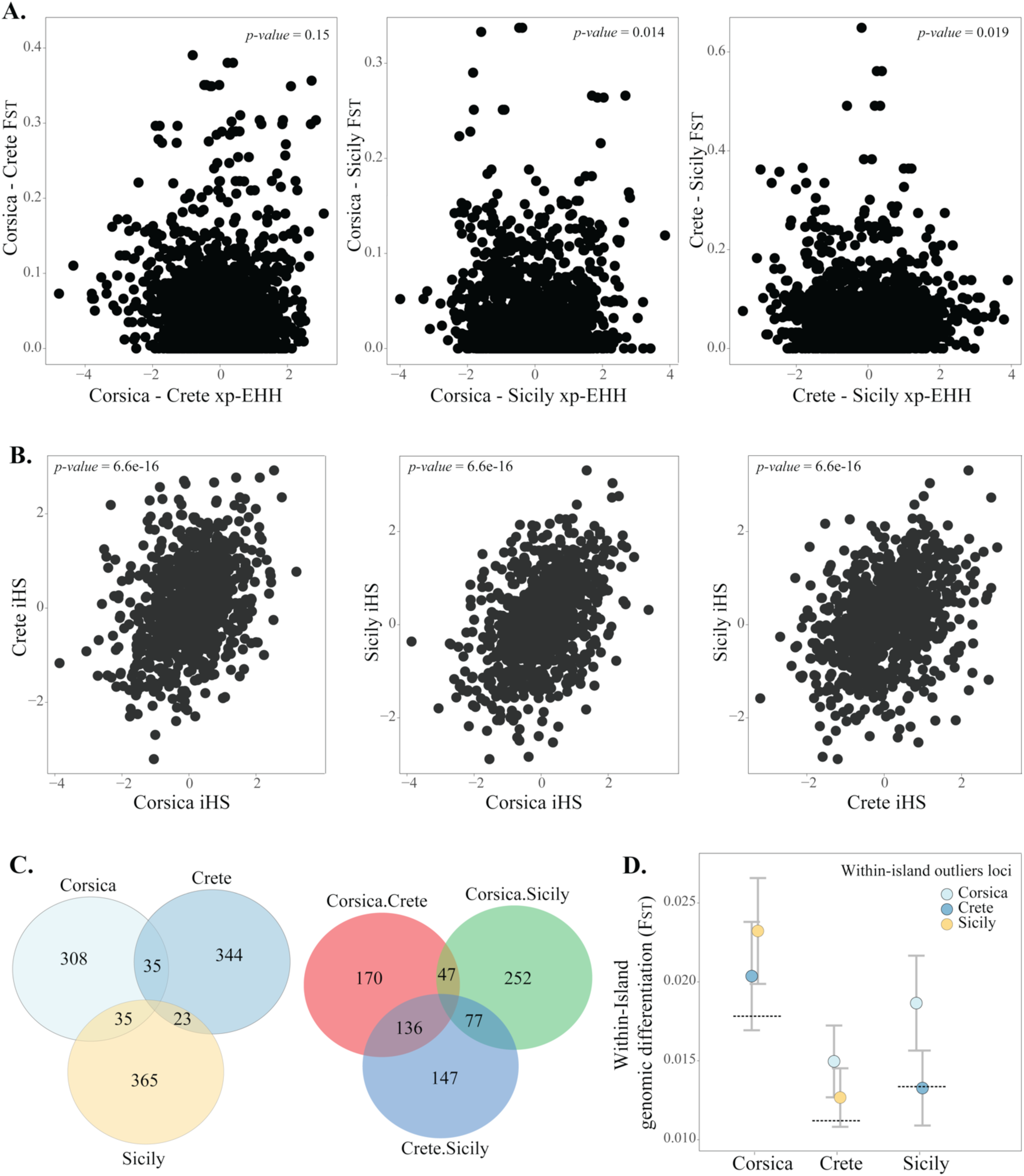
Effects of selection on genomic differentiation, similarity in selection pressures across islands and shared patterns of genomic differentiation. **A.** Relationship between divergent selection (Extended Haplotype Homozygosity statistic - XP-EHH) and genomic differentiation b*etween*-island. **B. Similarity in patterns of selection.** Correlations of *within*-island selection, measured as integrated haplotype homozygosity score (iHS) between islands. **C.** Shared outlier loci among populations *within*-islands (right) and *among*-islands (left). **D.** *F*_ST_ for outlier loci from each of the other islands within each island. Dashed lines represent the mean global within-island *F*_ST_ mean. Error bars denote 95% CI.

We also addressed how consistent selection is within and across islands. Patterns of *within*-island selection (iHS) showed a significant positive correlation in all pairwise comparisons between islands, with R^2^ ranging from 0.095 to 0.17 for iHS (Fig. 5B; Table S9C). This suggests shared patterns of selection across the genome among islands, potentially driven by similar selection pressures or genomic constraints reducing the availability of genomic variation. However, differentiation within island populations is not significantly correlated to selection (iHS) for Crete or Corsica (Fig. S2; Table S9B), although there is a weak significant correlation for Sicily (R^2^=0.005, *P*=0.01; Fig. S2; Table S9B). Interestingly, mean Tajima’s D differs considerably among islands, with Sicilian and Corsican populations exhibiting negative estimates (range -0.25 to -0.048; Fig. S3; Table S3). In contrast, populations on Crete exhibit higher values of Tajima’s D (-0.11 to 0.015; Fig. S3; Table S3).

### IV) Distribution of differentiation across the genome

We found that patterns of differentiation are significantly correlated between Sicily and Corsica (R^2^: 0.0066, *P* = 1.16e-10), but not between Corsica and Crete (R^2^: 5.8e-4, *P* = 0.16), or Sicily and Crete (R^2^: 1.96e-4, *P* = 0.81; Fig. S4). Hence, levels of differentiation are not correlated between all islands. Interestingly, we find that the outlier loci that are most differentiated within one island occur more frequently among other island outliers than expected by chance in two out of three comparisons (Fig. 5C; Table S10). A total of 9.3% of the outliers within Corsica overlap with those from Crete (Chi-squared: 7.18, *P:* 0.007) and a similar percentage is found in within-Sicily *F*_ST_ outliers (9.3%; Chi-squared: 6.80, *P:* 0.009; Table S10). However, Crete and Sicily outliers do not significantly overlap (Chi-squared: 0.09, *P:* 0.767). We also tested whether individual island outliers also have a higher mean distribution of *F*_ST_ values within other islands. We find that outliers from Sicily have a higher *F*_ST_ than the mean among Corsican populations (t = -3.082, df = 444.7, *P* = 0.002; Fig. 5D) and among populations from Crete (t = -1.997, df = 457.9, *P* = 0.046). Similarly, Corsica *F*_ST_ outlier loci have higher *F*_ST_ within Sicily (t = -3.393, df = 392.2, *P* = 7.6e-4) and within Crete (t = -3.586, df = 385.1, *P* = 3.8e-4). However, outliers from Crete do not have higher *F*_ST_ values than expected by chance within Corsica (t = -1.403, df = 400.3, *P* = 0.161), or Sicily (t = 0.078, df = 425.1, *P* = 0.938; Fig. 5D).

Pair-wise correlations between differentiation within- and among islands suggest that while differentiation within Corsica is correlated to differentiation between Corsica and Sicily (R^2^: 0.0025, *P*: 0.013) and Crete and Sicily (R^2^: 0.0025, *P*: 0.026; Fig. S5), none of the other comparisons are significant. Consistent with this pattern, the proportion of the most differentiated loci within Corsica that overlap with the most differentiated loci between Corsica-Sicily (9.8%) and between Crete-Sicily (7.4%) are also higher than expected by chance (Chi-square tests: *P*: 8.1e-05 and *P*: 0.04, respectively; Table S11). Moreover, we also find that a higher proportion (10.2%) of loci that are most differentiated within Crete than expected by chance are among those most differentiated between Crete and Sicily (*P*: 4.3e-06, Table S11).

### V) Patterns of local genomic differentiation in relation to parental contributions to the **genome**

The factor that best predicts the probability that a SNP is an outlier or not within islands is the extent of differentiation to the house sparrow (Logistic regression estimate: -1.401e+00, *P:* 0.036; Table 2). However, when evaluating factors that may affect the within-island differentiation (*F*_ST_), using a GLM, the differentiation to the house sparrow was found to be non-significant (Table S7). Neither the extent of differentiation from the Spanish sparrow, parental differentiation, recombination rate, nor hybrid index (HI) contributed significantly to differentiation, in both models, the logistic regression or the GLM (Table 2; Table S7). In separate logistic regressions for each island, none of the studied factors significantly affected the probability of being an outlier (Table S6), potentially because of reduced statistical power. While in separate GLMs per island (Table S8), general distribution of differentiation (*F*_ST_) between Corsican populations is explained by the differentiation to the Spanish sparrow (Estimate=1.166e-02, *P*=0.036) and within-Sicily differentiation is significantly explained by recombination rate (Estimate=-3.475e-03, *P*=9.3e-3; Table S8).

Taking advantage of the differences in parental contribution among islands, we addressed whether differentiation within islands was correlated to differentiation between the focal island population and the minor-ancestry parent species. For Corsica and Sicily, there is a significant correlation between within-island differentiation and that against the house and Spanish sparrow, respectively (Corsica: R^2^ = 0.002, *P*: 3.58e-4; Sicily R^2^ = 0.001, *P:* 0.022; Fig. 6A) but not to the alternative parent species in either case (Fig. S6B). Population differentiation within Crete was not correlated to differentiation to any of the parental species (Fig. 6A; Fig. S6B). Similarly, loci involved in high population differentiation *within*-islands (1% *F*_ST_ outliers) were more differentiated to the minor-ancestry parent than the genome wide neutral expectation (Fig. 6B). Outliers from Corsica and Crete present higher genomic differentiation to the Spanish sparrow, parent contributing the least ancestry, than the overall genome-wide average. However, the same pattern is not found in Sicily, where the house sparrow is the minor-ancestry parent, *within*-island outliers do not show higher differentiation than the genome-wide average (Fig. 6B). Also, outliers from Corsica were significantly more differentiated from Spanish- than house sparrow (t = -6.22, df = 519.4, *P* = 1.01e-09), as were outliers from Crete (t = -2.96, df = 668.2, *P* = 3.17e-3). And, outliers within Sicily, where the Spanish sparrow is the major-ancestry parent, are more differentiated from house sparrow than from Spanish sparrow (t = 3.76, df = 679.8, *P* = 1.81e-4; Fig. 6B). While divergence between the parental species (house-Spanish *F*_ST_) is weakly correlated to population differentiation within Corsica (R^2^ =3.4e-3, P=5.34e-5) and it is not significantly correlated to differentiation within the other islands (Fig. S6A). Similarly, within Corsica, outlier loci also have higher parental differentiation than expected by chance (t = 2.15, *P* = 0.033; Fig. 6B), but this does not hold true for Crete (t = 1.852, *P* = 0.065) or Sicily (t = -0.811, *P* = 0.42). Differentiation *within* Corsica and *within* Sicily was not correlated to differentiation *within* either of the parental species (Fig. S7) and differentiation *within* the parental species was not higher than expected by chance in outlier loci for these two islands (Fig. S8). Differentiation *within* Crete, and highly differentiated loci, was however, negatively correlated to differentiation *within* the house sparrow (Fig. S7; Fig. S8). These results suggest that overall, outlier loci for *within*-island differentiation are not involved in the genomic differentiation among populations within each parent species.

**Figure 6.**
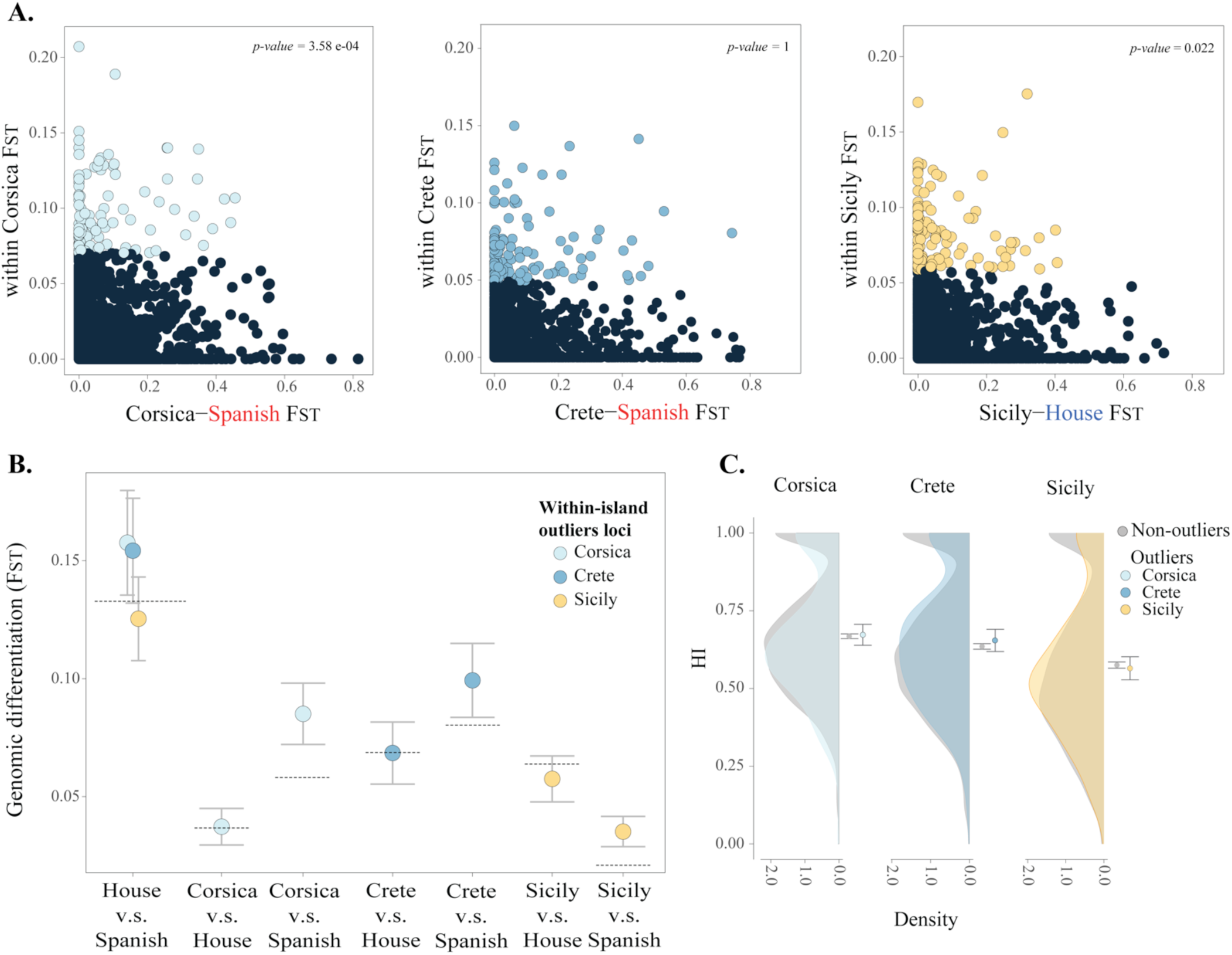
Effects of divergence from parental species and minor –major parent ancestry on differentiation. **A.** Correlations of between populations *within*-island *F*_ST_ and genomic differentiation from the minor-ancestry parent (island v.s minor-parent *F*_ST_). **B.** Between parents divergence (house-Spanish *F*_ST_) and island-parent *F*_ST_ for *within*-island outlier loci compared to the genome wide average (dashed line). **C.** Density of hybrid index (HI) for within-island *F*_ST_ outliers compared to the genome wide distribution. Means and 95% confident intervals of outlier and non-outlier loci depicted.

Finally, we assessed the effect that ancestry divergence can have on the genomic differentiation within islands. The distribution of the hybrid index (HI) does not differ between the *within*-island *F*_ST_ outliers and the non-outlier loci for any of the islands (t-test of t=0.27, 1.04 and -0.53, for Corsica, Crete and Sicily, respectively. P>0.05 for all islands; Fig. 6C). A post hoc correlation analysis shows no evidence to support that the overall genomic differentiation within islands is largely explained by ancestry (R^2^ varying between 5.1e-4 and 1.8e-5, with P>0.05 for all islands; Fig. S9A). We further tested if the HI affected the probability of a locus to be highly differentiated within islands (1% *F*_ST_ outlier loci; Fig. S9B). Examining outliers with extreme values of HI only, we found that outliers within Sicily more frequently have excess Spanish ancestry (mean frequency proportion based on 10000 random draws = 0.31), whereas Corsica and Crete have outliers with excess house ancestry (mean frequency proportion based 10000 random draws = 0.804 and 0.799, respectively; Fig. S8B). Even though comparison with proportions from a similar resampling analysis for non-outlier loci shows that distributions differ (Corsica: t = 27.175, Crete: t = -30.846, Sicily: t = -4.369, all *P<*1.25e-05), the ancestry pattern of outliers is generally very similar to that of the general genomic background (Fig. S9B).

## DISCUSSION

While evidence for a creative role of hybridization in evolution is piling up, little is known about how the genomes of hybrid taxa can differentiate in response to local selection pressures. Investigating the factors that explain how hybrid taxa can differentiate within lineages of the Italian sparrow, we find high within-island differentiation at relatively short distances in the light of the dispersal ability of the species [48]. A discriminant function analysis classifies 75-95% of the individuals to the correct population within islands. This local differentiation suggests that there is potential for population structure and adaptive divergence within this hybrid species. However, there is also pronounced between-island differentiation, approximately five times higher than within-island differentiation. Interestingly, genomic differentiation and a measure of between- islands selective sweeps are significantly correlated in two out of three between-island-comparisons, supporting a role for natural selection in shaping the differentiation of the hybrid genomes of Italian sparrows in geographically isolated islands. However, contrary to our expectation of positive correlations, the negative nature of the relationship do not support the scenario where highly differentiated parental alleles are involved in divergent natural selection facilitating genomic differentiation between islands. Genomic differentiation within island is poorly explained by signatures of selective sweeps, with the exception of Sicily, where patterns of selection are weakly correlated with local genomic differentiation. In this species, we know from previous findings that local differences in beak shape among island populations is driven by climate and diet [43] and adaptive differentiation in mainland Italy is also driven by climatic variables [32, 42]. This is also consistent with the large body of work suggesting that hybridization enables adaptive variation across a range of taxa [2,49,50,51].

Interestingly, there is some evidence that the same genomic regions repeatedly are involved in population divergence. Differentiation within Corsica is significantly correlated to that within Sicily, but differentiation within Crete is not correlated with that of the other islands. Although our analyses may lack statistical power to detect such correlations, this could also be due to the contribution of *P. domesticus biblicus,* a house sparrow subspecies distributed across the Middle- East, to the population on Crete (Ravinet *et al*., in prep). This introgression may also have contributed to Crete forming a third cluster in the Admixture analysis. In addition, Corsica also shares a higher proportion of the most strongly differentiated outliers than expected by chance with both Crete and Sicily, while the proportion of outliers shared between Crete and Sicily is not higher than expected by chance. These results may, to some degree, support the hypothesis that loci linked to local genomic differentiation may be limited to specific genomic regions and are reused across hybrid lineages.

Interestingly, patterns of selection are consistent across populations and correlated between islands. Hence, the findings of some degree of shared differentiation could partially be explained by a similar selection landscape for all populations of this hybrid taxon or by specific allelic combinations available to selection. Earlier work has shown that the same genetic composition as in the wild ancestor repeatedly arise in lab-crosses of *Helianthus* sunflowers [36]. It remains to be investigated to what extent the similarity in selection landscapes is caused by historically shared selection in the ancestral populations of the parental species (Ravinet *et al*. in prep), selection for a functional admixed genome [22, 29], stabilizing selection, linked to human commensalism [52] or parallel selection linked to adaptation to insularity. For instance, such shared ancestral selection landscape could lead to bias in which parental alleles are retained or more prone to be lost or selected against (i.e. the Spanish sparrow being considered a non-human commensal in most of its range, compared to the house sparrow).

A major finding is the limited evidence of genome structure shaping local differentiation. Variation in the underlying recombination rate landscape may mould the landscape of differentiation [53], and has been shown to affect e.g. the genomic differentiation generating correlated patterns of differentiation in divergent populations of mice, rabbits [40] and flycatchers [37]. In admixed species, selection against minor parent ancestry has been hypothesised to generate patterns of strong correlation between measures of introgression and recombination rate [20,38,41]. This type of selection might be expected to reduce the genetic variation available for differentiation among hybrid populations. As only the latter process is hybrid specific, a decoupling of the correlation between recombination rate and differentiation present among the parent species is expected if purging of minor parent alleles is important during stabilization of hybrid genomes (Fig. 1A). We find that within island divergence is significantly less affected by recombination rate than divergence between the parent species. This could suggest that selection against incompatible minor parent alleles in low recombination regions reduces the potential for differentiation within the hybrid species in these regions. Overall, very little of the divergence between populations within islands is explained by recombination rate, despite its weak correlation with differentiation in Sicily. Furthermore, recombination rate overall did not significantly improve models explaining within island differentiation.

Differentiation between the parent species could potentially affect the diversity available for adaptation (Fig. 1C). Low parental divergence is expected to reduce the number of segregating alleles for selection to act on, consistent with findings that hybrids from more divergent parent species are morphologically more novel [54, 55]. However, genomic regions of high divergence between parental species can also harbour potential genomic incompatibilities in the hybrid taxon. This could generate a negative relationship between the genomic differentiation in the hybrid populations and highly divergent parental loci as the hybrid can only fix ancestry from one of the parent species (Fig. 1C). Indeed, we find that differentiation between the parent species explains neither the degree of differentiation within islands, nor improves the fit of the models evaluating *within*-island differentiation (logistic and GLM-models). Interestingly, a similar pattern has previously been found in the populations of the Italian sparrow in mainland Italy [32], suggesting that constraints might have played an important role during the stabilization of the hybrid genome, limiting genomic variation. The repeatability of this result across the range of the Italian sparrow also suggests that hybridization may indeed lead to predictable patterns of genomic variation long after genome stabilization and hybrid species formation. We find that differentiation from the house sparrow is the factor that best explains the probability of being a highly differentiated locus within islands. An additional source of variation in hybrid species could stem from variants that segregate within the individual parental species. However, we did not find any compelling evidence that within-parent differentiation predicts differentiation within the hybrid species.

Whether ancestry is a determining factor on how genomic differentiation is distributed in the hybrid genome is not easily disentangled. The divergence in ancestry proportion from the minor – major parent in the Italian sparrow [22] enables us to test whether the differences in ancestry has affected population differentiation. Purging of genomic incompatibilities of the minor parental ancestry blocks plays an important role in determining the genetic variation in the Italian sparrow [27, 47]. A range of studies has suggested that the probability of retaining neutral ancestry is higher in genome regions with a high recombination rate [20,38,41,56]. To address if minor parent ancestry, in spite of selection against incompatibilities, could be involved in adaptation within the Italian sparrow, we investigated whether minor parent ancestry was important for differentiation within the Italian sparrow. We did not find a clear effect of ancestry in population differentiation within islands; nevertheless, overall highly differentiated loci are not found more frequently in minor-parental ancestry blocks. However, we found significant correlations between local differentiation and the differentiation to the minor-ancestry-parent for two out of three islands. Overall *F*_ST_ outliers are also more differentiated from the minor-parental blocks than expected based on genome-wide levels of differentiation. This suggests that loci linked to local differentiation *within*-islands are also involved in divergence to the minor-ancestry parent. Recombination rate can determine how ancestry is distributed across the hybrid genome [20, 38] and this may affect the effect ancestry has on genomic differentiation. The probability for minor-ancestry blocks to rapidly decouple from potential incompatibilities with the major-parent genetic background, increases with recombination rate [20], which in turn can have an effect on how easily regions with minor-ancestry in high/low recombination areas are able to harbour genomic differentiation among populations.

Interestingly, the probability of being among the 1% most strongly differentiated loci is best explained by how differentiated a given island population is to the house sparrow. Neither recombination rate, the ancestry of the region, nor the differentiation to the Spanish sparrow significantly affected the degree of differentiation or the probability that the locus was an outlier. This is an interesting finding as [22] previously also found a skew towards house sparrow ancestry in loci consistently inherited from one parent species, particularly identifying an enrichment of mito-nuclear loci and loci involved in DNA-repair. Potentially, this could be indicative of some constraints on differentiation from the house sparrow, as most Italian sparrows are fixed for house sparrow mitochondrial haplotypes [5, 22].

### Conclusion

Taken together, our findings suggest that hybrid lineages may use their elevated levels of variation for local adaptation, with differentiation among islands being significantly correlated to differences in signatures of selective sweeps in two out of three comparisons. Moreover, the findings of correlated differentiation among islands as well as similar selection pressures within islands may suggest that similarity in selection pressures and/or constraints can contribute to parallelism in genome evolution in the hybrid Italian sparrow. Interestingly, we find that the negative relationship between recombination rate and differentiation expected due to linked selection being stronger in low recombination regions, was significantly stronger in the parent-parent comparison than within the three hybrid lineages. This would be consistent with a lower differentiation in low recombination regions within the hybrid lineages, as expected if purging of minor parent alleles reduces the variation available for divergence. However, a logistic model revealed that differentiation to the house sparrow is the overall best predictor of the probability of outlier status. Jointly, this suggests that selection interacts with constraints linked to admixture and past hybridization during the stabilization of hybrid genomes. To further understand how constraints and selection shape hybrid genomes, one interesting future line of inquiry would be investigating whether alleles inherited from the parent contributing least to the genome in genomic regions involved in population differentiation are more likely to occur in high recombination rate regions.

## MATERIALS AND METHODS

The Italian sparrow originated from hybridization between house and Spanish sparrow, likely during the spread of the commensal house sparrow to Europe in the wake of the introduction of agriculture [52, 57]. The parental species diverged approximately 0.68 million years ago [52]. In addition to the distribution on the Italian peninsula, Italian sparrow populations are also found on some Mediterranean islands. These insular populations have strongly differentiated genomes, with different contributions from each parent species [22], and exhibit phenotypic divergence with island specific beak shape matching local temperature and diet [43]. Furthermore, the island populations are evolutionarily independent and are hypothesized to have arisen from individual hybridization events.

### Sampling and sequencing

Three populations of Italians sparrows were sampled from each of the islands Sicily (n=76), Crete (n=77) and Corsica (n=70) in March-June 2013 (Fig. 2A, Table S1). On each island we sampled individuals from three geographically separated populations (Fig. 2B, Fig. 3C). Population sample size varied between 16 and 30 (Table S1). We sampled reference house sparrow parent populations from Norway (n=11), and Spanish sparrows from Kazakhstan (n=10). To increase the number of sampled individuals from the parent species for analyses that work better with approximately equal sample sizes of all taxa, we added house sparrow samples from Switzerland (n=17) and France (n=18) and Spanish sparrow samples from the Gargano peninsula (n=14) and Spain (n=23); (Table S1). All birds were caught using mist nests, and blood was sampled from the brachial vein and stored in Queens lysis buffer. All necessary permits were obtained from relevant local authorities prior to sampling. DNA was extracted using the Qiagen DNeasy Blood and Tissue Kit, (Qiagen N.V., Venlo, The Netherlands) and the product was stored in Qiagen’s buffer EB prior to sequencing. We used a RAD-tag approach, and library preparation, sequencing, demultiplex and removal of adapters were done by Ecogenics GmbH (Balgach, Switzerland; www.ecogenics.ch). Specifically, the restriction enzymes EcoRi and MseI were used for double digest restriction-site associated DNA sequencing (ddRAD).

### Data processing and variant calling

First, the quality of all RAD sequencing was checked using FASTQC [58]. Raw reads were filtered using the module process_radtag from the software Stacks [59]. Reads shorter than 73 base pairs were discarded as well as those with an uncalled base. To ensure high confidence-based calls, a Phred quality score of 20 (99.9% accuracy) was used as threshold across a sliding window fraction of 0.1 of the read length. We used BWA-MEM (v 0.7.8) [60] to map the reads to the house sparrow reference genome [5] using default parameters. We re-aligned indels realignment with GATKs (v 3.7) RealignerTargetCreator and IndelRealigner [61, 62] and called the variants using HaplotypeCaller [62]. For a detailed description of the variant pipeline, see [32]. We filtered SNPs using Vcftools v. 0.1.14 [63], setting the filter parameters to --max-missing 0.8, --minDP 10.00, -- minGQ 20.00 and --maf 0.02. We pruned linked sites using PLINK v. 1.9 [64] with 100 kb windows, 25 bp step and an R^2^ threshold of 0.1. VCF-files containing different set of individuals were generated to suit the different analyses (Table S2).

### I) Genomic differentiation within- and between islands

We tested the hypothesis that the degree of divergence is significantly higher between islands than within islands, reflecting long periods of independent evolution. To this end, we first illustrated the overall divergence between the islands and populations using a Principal component Analysis as implemented in glPca() in the R package ADEGENET 2.0 [65]. We also evaluated the level of clustering in the data through estimating the cross-validation error for K=1 to K=9, and estimated the probability of each individual belonging to these clusters using ADMIXTURE v.1.3.0 [66] . To illustrate the extent to which the divergence was aligned with the axis of parental differentiation, three parental populations of each species were included in these analyses, resulting in 316 individuals approximately equally distributed across the three species (Table S2). After filtering for LD, the remaining 2224 SNPs were included in the PCA and Admixture analyses. To further illustrate the degree of differentiation within islands, we also performed a Discriminant Analysis of Principal Components (DAPC) within each island to address to which extent the local populations can be correctly classified based on the available variation. To characterize potential variation in genomic diversity we also estimated nucleotide diversity in sliding windows of 100kb in size with 25-kb steps for each population using vcftools v. 0.1.14 [63]. For estimates of nucleotide diversity non-variant sites were retained, and we did not filter on minor allele frequency.

We investigated whether the differentiation was stronger between islands than within islands, using two approaches. First, we estimated global *F*_ST_ among populations within islands, as well as pair-wise *F*_ST_ among islands in 100kb windows using vcftools v. 0.1.14 [63]. The window size was selected as linkage disequilibrium in sparrows is known to decay within this distance [5]. We used a Monte Carlo permutation paired t-test to investigate if pairwise *F*_ST_-values were higher among- than within islands.

Second, we used an Analysis of Molecular Variance (AMOVA) to formally address what proportion of genetic variance is explained by differentiation among islands, among local populations within islands, within local populations and variation within individuals. We transformed the VCF to a genlight object, where levels of divergence were defined, using the ADEGENET R-package and ran an AMOVA with the poppr.amova() function from the POPPR R-package [67, 68]. We assessed significance by a randomization of population assignments using a Monte Carlo test with 1000 permutations implemented in the randtest() function from the ADE4 R-package [69]. See supporting methods for additional analyses assessing the sensitivity to missing data.

### II) The relationship between genomic differentiation and recombination rate

We examined the hypothesis that hybrid genome formation influences the association between differentiation and recombination rate. Our rationale was that if purging of minor ancestry is stronger in low recombination regions, this reduces diversity in these regions and therefore acts as a constraint on differentiation such that we expect greater differentiation in higher recombination regions where the effect of purging is weaker. Alternatively, if local selective sweeps play a more important role in shaping hybrid genomes, we would expect greater differentiation in low recombination regions [37, 40]. As the relative strength of these processes is unknown, we used the relationship between differentiation and recombination rate between the parent species as a null expectation, and tested if there was a deviation from this relationship in the direction expected from purging of minor parent ancestry in the hybrid populations compared to the parent species (see Fig. 1A). To this end, we tested for differences in the slopes of individual linear regressions of *F*_ST_ and recombination rate. We also evaluated a significant interaction between lineage combination (parent-parent vs. within island) and recombination rate on *F*_ST_ using independent linear model per island. We used recombination rate estimates from [5]. We also evaluated the significance of the relation between genomic differentiation and recombination rate within each island using Pearson’s correlation tests. We used Bonferroni corrected *P-*values to account for multiple comparisons.

### III) The concordance of patterns of selection and genomic differentiation

To address if elevated genomic differentiation is driven by strong divergent selection, we performed Bonferroni corrected Pearson’s correlations of *F*_ST_ between island pairs to their cross-population Extended Haplotype Homozygosity statistic (xp-EHH) [70], which is designed to compare haplotype lengths between populations (between islands in this case) in order to detect selective sweeps. We further investigated whether regions putatively under selection within-island are independent across islands and whether they coincide with areas of elevated differentiation. We performed pairwise Bonferroni corrected Pearson correlations between each island pair of the integrated haplotype homozygosity score (iHS) [71] developed for detecting positive selection within a population, in this case calculated within each island. Then, we tested if putative concordance in selection may result in correlated patterns of differentiation in islands, through investigating the correlation between iHS-scores and genomic differentiation for each island. We estimated long range haplotype statistics through phasing data with SHAPEIT/v2.r837 [72, 73] and converted the resulting VCF-file using the vcfR R-package [74]. We then used the functions data2haplohh(), ihh2ihs() and ies2xpehh() from the rehh R-package [75, 76] to prepare the data, estimate the integrated haplotype homozygosity score (iHS) and estimate Extended Haplotype Homozygosity (XP-EHH), respectively.

### IV) Distribution of differentiation across the genome

To test if the differentiation landscape between populations within islands is correlated to that within other islands and between islands, as would be expected if differentiation is affected by the underlying recombination rate landscape and constraints or similar selection pressures acting on the populations, we performed pairwise Pearson’s correlation tests on *F*_ST_ estimates. We tested if global *F*_ST_ estimates within one island were significantly correlated to these within another island, as well as if between-island differentiation was significantly correlated to global *F*_ST_ within any of the islands using a resampling approach and Bonferroni corrections for multiple testing.

In addition, we investigated to what extent the same loci were among the most strongly differentiated on different islands. We estimated the proportions of the 1% most differentiated loci that were shared between each island pair. We then investigated if this proportion of shared *F*_ST_ outliers was higher than expected by chance using a series of χ^2^-test for each pair-wise comparison, applying Bonferroni corrections for multiple testing.

### V) Patterns of local genomic differentiation in relation to parental contributions to the genome

Multiple factors may affect which loci are free to vary within the Italian sparrow. For example, variation in parental contributions to the genomes of the different island populations, the level of differentiation between the parent species across the genome and the recombination rate. We tested to what extent these factors explain the patterns of within-island differentiation by performing a generalized linear model (GLM) using within-island *F*_ST_ as the response variable: *F*_ST_ = ancestry proportion (hybrid index) + recombination rate + island + island to house sparrow differentiation (*F*_ST_) + island to Spanish sparrow differentiation (*F*_ST_) + parental differentiation (house-Spanish *F*_ST_). We also evaluated how these factors affected the probability of a locus to belong to the 1% most differentiated loci within an island using a similar model with a logistic regression where the response variable was the Pr(outlier). In addition, we performed logistic regressions and GLM individually for each island, excluding the island term. As post hoc tests, we examined Bonferroni corrected Pearson correlations of within-island differentiation against differentiation of the island to each of the parental taxa as well as between the parent species. We also assessed whether highly differentiated loci found in the Italian sparrow are also involved in the genomic differentiation among populations within each parent species (within-house *F*_ST_ and within-Spanish *F*_ST_).

Finally, to address if variation in minor parent ancestry affects within-island differentiation, we tested the correlation between genomic differentiation and hybrid index (HI) reflecting the relative contribution of each parent species. We also tested if highly differentiated loci were found in blocks with high allele frequencies from major- (less than 35% minor parent alleles) or minor parent ancestry (less than 35% major parent alleles) more frequently than expected by chance by comparing the confidence intervals from 10000 resamplings of 8 outlier loci to the value for the entire *F*_ST_-distribution to assess significance. The same analysis was run for the distribution of non-outlier loci to assess whether the outliers diverge from the neutral expectations. Hybrid index (HI) was estimated using whole genome data from [22] for Crete, Corsica and Sicily, including parental species from [52] and [5]. We phased data using SHAPEIT/v2.r837 [72, 73] and estimated ancestry using LOTER [77] and translated this into a per locus hybrid index, where 0 corresponds to only Spanish ancestry and 1 corresponds to pure house sparrow ancestry. We estimated the hybrid index separately for each island.

## ETHICS STATEMENT

All relevant sampling permits were obtained from the regional authorities and handling of birds was conducted according to their guidelines. (Museum National d’Histoire Naturelle, Centre de Recherches sur la Biologie de Populations d’Oiseaux, Paris (France), Institute for Environmental Protection and Research – ISPRA (Italy), Consejería de Industria, Energía y Medio Ambiente (Spain), Norwegian Food Safety Authority (Norway), and Bundesamt für Umwelt BAFU, Abteilung (Switzerland)) and Ministry of Education and Science (Republic of Kazakhstan).

## ACKNOWLEDGEMENTS

We thank Jo S. Hermansen and Maria Tesaker for help with fieldwork. This research was funded by a Wenner-Gren fellowship to A.R., a Norwegian Research Council grant to G-P.S. and A.R., and a Marie-Curie Fellowship and a University of Oslo, Faculty of Sciences grant to F.E.

## SUPPORTING INFORMATION

**Table S1. Sampled individuals from the parent species (the house and Spanish sparrows) and the Italian sparrow.**

**Table S2. VCF files.**

**Table S3. Per-island population genomic statistics.** Mean values of π and within-island genomic differentiation (F_ST_). t-test for pairwise comparison between genome wide within island F_ST_. Mean values of Tajima’s D per population within each island.

**Table S4. Intercept, slope and confidence intervals of the slope of individual linear regression of *within*-island genomic differentiation and recombination rate as well as parent-parent differentiation and recombination rate.**

**Table S5. Evaluating the effect that the interaction between recombination rate and the type of comparison (parental differentiation (house-Spanish), which is the null model, and within-island differentiation) has on genomic differentiation (*F*_ST_).** Individual linear models per island were run to test if there is a significant interaction between recombination rate and comparison, as expected if the relationship between recombination rate and differentiation differs between parent species and the hybrid Italian sparrow (Fig. 1A).

**Table S6. Logistic regressions per island, on the probability of being a local F_ST_ outlier within island.**

**Table S7. Generalized linear model on within-island F_ST_.**

**Table S8. Generalized linear models per island on within-island F_ST_.**

**Table S9. Concordance of within-island selection (iHS) and divergent selection (xp-EHH with genomic differentiation.**

**Table S10. Number and percentage of within-island outlier loci shared between islands.** Chi-squared denote tests for overrepresentation compared to the genome wide average.

**Table S11. Number and percentage of within-island outlier loci identical to between-island outliers.** Chi-squared denote tests for overrepresentation compared to the genome wide average

**Figure S1. AMOVA significance - Randomization via permutation**. Monte Carlo test with 1000 permutations implemented in the randtest() function from the ADE4 R-package to evaluate significance. Black line denotes the observed values of Sigma (Variance in each hierarchical level).

**Figure S2. Concordance of patterns of selection and genomic differentiation**. Correlations of the integrated haplotype homozygosity score (iHS) and genomic differentiation (within-island FST).

**Figure S3. Distribution of Tajima’s D per population in each island**.

**FigureS4. Correlation of within-islands differentiation across the three Mediterranean islands**. Bonferroni corrections of the p-values are reported.

**FigureS5. Correlation of within-islands differentiation v.s. between-islands divergence**. Adjusted p-values after resampling and Bonferroni corrections.

**FigureS6. Correlation of within-islands differentiation and the parental species**. Adjusted p-values after Bonferroni corrections.

**Figure S7. Correlations of within-island differentiation and within-parent differentiation (within-house or and within-Spanish sparrow)**.

**Figure S8. A**. Intraspecific genomic differentiation in the parental species for the within-island F_ST_ outlier loci. Dash lines represent the within-parent F_ST_ global mean. **B.** t-tests evaluating weather within-island outlier loci present higher/lower values than expected by chance in the within-parent differentiation

**Figure S9. A.** Relation between within-island F_ST_ and ancestry (hybrid index - HI). Results of linear regression reported. Dashed lines depict the 1% outliers threshold. **B.** Frequency proportion of outlier loci found in regions of mainly house ancestry (0.65<HI) and mainly Spanish ancestry (HI<0.35) (minor-major parental ancestry). Distribution of 10.000 random resampling draws of 8 outlier loci.

## DATA ACCESSIBILITY

Genomic data will be deposit at the NCBI Sequence Read Archive or at the European Nucleotide Archive (ENA) at http://doi.org/[doi], reference number [reference number]. Other data will be deposited in the Dryad Digital Repository at [accession numbers or DOI will be provided once the data is uploaded]. Data will be uploaded upon acceptance of the manuscript.

## AUTHORS’ CONTRIBUTIONS

A.R. conceived and designed the study with advice from G- P.S. and F.E.; A.R., F.E. and G-P.S. collected field data; A.R. and A.C. wrote the manuscript; A.C. conducted the laboratory work; A.C. developed the RAD-data analysis pipeline with input from M.R.; A.C. and A.R. designed the analyses; A.C. analysed the data with contributions from F.E., M.R. and A.R.; all authors discussed the results and commented on earlier drafts of the manuscript.

